# Anubis: a multi-level authentication scale for ancient proteins using random forest classification

**DOI:** 10.1101/2024.11.15.623824

**Authors:** Yun Chiang, Bharath Anila Bhuvanendran Nair, Max Emil Ermter Ramsøe, Tina Ravnsborg, Ole Nørregaard Jensen, Matthew James Collins

## Abstract

Deamidation is a time-dependent, post-translational modification with wide applications in clinical proteomics and palaeoproteomics as ageing indicators and authentication markers. Despite its significance, the use of deamidation for authenticating ancient proteins remains controversial, primarily due to variability in deamidation kinetics and context-specific protein preservation. We propose `Anubis`, an open-source, Python-based pipeline to evaluate relative deamidation patterns and authenticate ancient proteins. `Anubis` leverages random forest classification based on a range of tandem mass spectrometry-based features and physico-chemical characteristics. We use a combination of ancient and experimentally degraded beta-lactoglobulin to demonstrate that `Anubis` can authenticate a putative ancient dairy protein using position-specific deamidation patterns, relative deamidation (asparagine vs. glutamine), and overall modification patterns using trypsin processed in the same batch as a baseline. This multi-level approach addresses the complexities of ancient protein deamidation and paves the way for automated, reliable authentication across different archaeological substrates.

## Introduction

Spontaneous, non-enzymatic, and time-dependent deamidation is a significant post-translational modification (PTM) for long-lived proteins, for example, eye crystallins, collagens, and histones (basic proteins with abundant lysine and arginine residues) ^1–5^. It affects protein stability and aggregation, associated with the development of cataracts, Alzheimer’s, Parkinson’s, and neurodegenerative diseases ^6–10^. Hence, the accumulation of deamidated proteins have been widely used as ageing indicators and disease biomarkers.

Despite the biological and chemical importance of deamidation, detailed deamidation pathways are still poorly understood. Generally, asparagine deamidates via a ring intermediate (five-ringed succinimide) under neutral or basic conditions, in which the carbonyl carbon of an asparagine side chain is attacked by the nucleophilic backbone nitrogen of the next (n+1) amino acid ^11–14^. This succinimide intermediate is susceptible to hydrolysis and isomerisation, resulting in the formation of aspartic and iso-aspartic acids. Alternatively, direct side-chain hydrolysis leads to the formation of aspartic acid and the release of ammonia. While glutamine deamidation is also facilitated by a glutarimide intermediate or side-chain hydrolysis, the formation of a six-ringed glutarimide and side-chain hydrolysis are both slow processes with high activation energies ^15,16^. Since deamidation is a multi-step and multifactorial modification, actual deamidation kinetics are mediated by a range of chemical, structural, and environmental factors, including neighbouring amino acid sequences, structural conformations, solvent accessibility, pH, and temperature ^17–19^.

Given the time dependence and ageing-relevance of glutamine deamidation, it has been proposed as an authentication marker for ancient proteins ^20–23^. The underlying assumption is that authentic, endogenous ancient proteins would exhibit more advanced levels of deamidation than modern contaminants. However, given the context of ancient protein preservation, it is noted that the lack of deamidation is indicative of limited degradation, rather than contamination ^24^. Moreover, it has been highlighted that there is no clear relationship between time and deamidation in archaeological samples, and that deamidation is not a reliable authentication marker due to varying deamidation kinetics ^25^.

While the use of glutamine deamidation as an authentication tool remains inconclusive in palaeoproteomics, proper authentication is essential for research rigour and the confident identification of ancient proteins. Despite extreme cold or arid burial conditions ^26,27^, even the best-preserved ancient proteins are damaged, fragmented, and often low in abundance, easily masked by common and highly-abundant contaminants, including but not limited to modern humans. Multiple sources of contamination could also be introduced during archaeological excavations, handling, transportation, museum storage, and laboratory analysis ^28^. The history of ancient DNA (aDNA) research has demonstrated that the lack of authentication resulted in biases, misinterpretations, and controversies ^29–31^.

Despite the importance of ancient protein authentication, there are limited tools to automate this process for tandem mass spectrometry (MS2) data. Deamidation introduces a positive mass shift (+0.984016 Da, monoisotopic mass), and tandem mass spectrometry allows for the localisation of deamidation sites based on shifting fragment ion masses ^32^. One of the most commonly used, MS2-based tools is `DeamiDATE`, which was designed for `MaxQuant` (version 1.6.2.6) outputs ^23^; however, column names and formats were hard-coded and have not been updated for latest iterations (version 2.6.5.0 as of 2024). Additionally, `DeamiDATE` was executed by Python 2x, a sunsetted version with no further maintenance and support.

We propose `Anubis`, a multi-level and cloud-ready authentication system for ancient proteins and complex mixtures. Anubis was the Egyptian deity who weighed the hearts of the dead on a scale against the feather of Ma’at (the goddess of justice) ^33^. Similarly, the `Anubis` authentication pipeline weighs putative ancient proteins against common contaminants based on relative deamidation patterns at position-specific and protein levels. `Anubis` leverages a random forest (RF) classification model to incorporate 13 physio-chemical and MS2-based features to predict a deamidated peptide with probability. We focused on an RF model since it can handle a wide range of non-parametric features, with ensemble learning tolerating data noise and outlier events with high accuracy ^34,35^. We also include position-specific and protein-level deamidation comparisons to capture and mitigate deamidation variability. Anubis is open-source, modular, lightweight, and ready to be run in any Jupyter Notebook environments, especially Google Collaboratory (Colab).

We selected beta-lactoglobulin (BLG) for `Anubis` re-analysis, since it is the most abundant dietary protein recovered in the archaeological record ^36^. BLG has been associated with prehistoric sites when lactase persistence was absent ^37–43^, and it contains genus-specific sequences to distinguish *Bos* (cows) from *Caprinae* (sheep/goats) ^44^. Indeed, BLG is a low molecular weight (c. 18,300 Da) protein with a stable structure (a beta-barrel) ^45^, which may contribute to its long-term survivability. It is crucial to achieve the confident identification of ancient BLG, since it sheds light on palaeodiets, animal management, and potentially the co-development of dairy consumption and lactase persistence genes.

## Results and Discussion

### Evaluating the RF classification model

We built a binary RF classification model to predict whether a BLG peptide is deamidated using `Scikit-learn` and 13 physico-chemical, MS2-based features (Methods Table 1). The selection of these physicochemical indicators is consistent with previous RF models developed for monoclonal antibodies and drug therapeutics ^46,47^. We included MS2 data, since deamidation results in a mass shift and a change of retention time. MS2 quality characteristics, including MS2 intensity, the percentage of matched ion series, precursor mass difference, and false discovery rates (FDRs), were incorporated to see if ionisation efficiency, search engine scoring, and statistical controls would affect deamidation detection.

Given the accuracy paradox of classification modelling, accuracy (the chance of a model making a correct prediction) is an intuitively important concept, but not a reliable evaluation metric, depending on the balance of datasets ^48^. For example, without any data cleaning, we had 7,416 deamidated (class 1) and 20,249 non-deamidated (class 0) BLG PSMs, meaning that 73.22% accuracy is achievable if a model simply predicts all PSMs are non-deamidated. To mitigate this, we randomly selected 5,000 deamidated and an equal number of non-deamidated PSMs to achieve a balanced 50:50 class ratio. We also employed a range of evaluation metrics, including precision, recall, F1 scoring (a weighted mean of accuracy and precision), a confusion matrix (true positives vs true negatives), and Receiver Operating Characteristic (ROC) analysis (the ability of the model to correctly distinguish between classes).

We fine-tuned hyperparameters to optimise the RF model using a combination of randomised and grid searches (Methods). We randomly separated the BLG PSMs (n=10000) into 80% training, and 20% testing using `Scikit-learn`. The optimised RF model has high precision (91.00 %), recall (93.88 %), and F1 scoring (92.42%). This is supported by the confusion matrix with dominant true positive (93.88 %) and true negative (90.48 %) matches (Fig. 1a). For the ROC analysis, we included five runs of cross-validation to iterate through all BLG PSMs. The mean area under the ROC curve (AUC) is 0.9738, representing a 97.38% probability that the model correctly labels random positive and negative classes (Fig. 1b). Overall, the model is effective at predicting deamidated BLG PSMs without data loss, and can handle unseen data without issues. This is crucial in palaeoproteomics, since protein preservation tends to be context-specific.

**Fig. 1.**
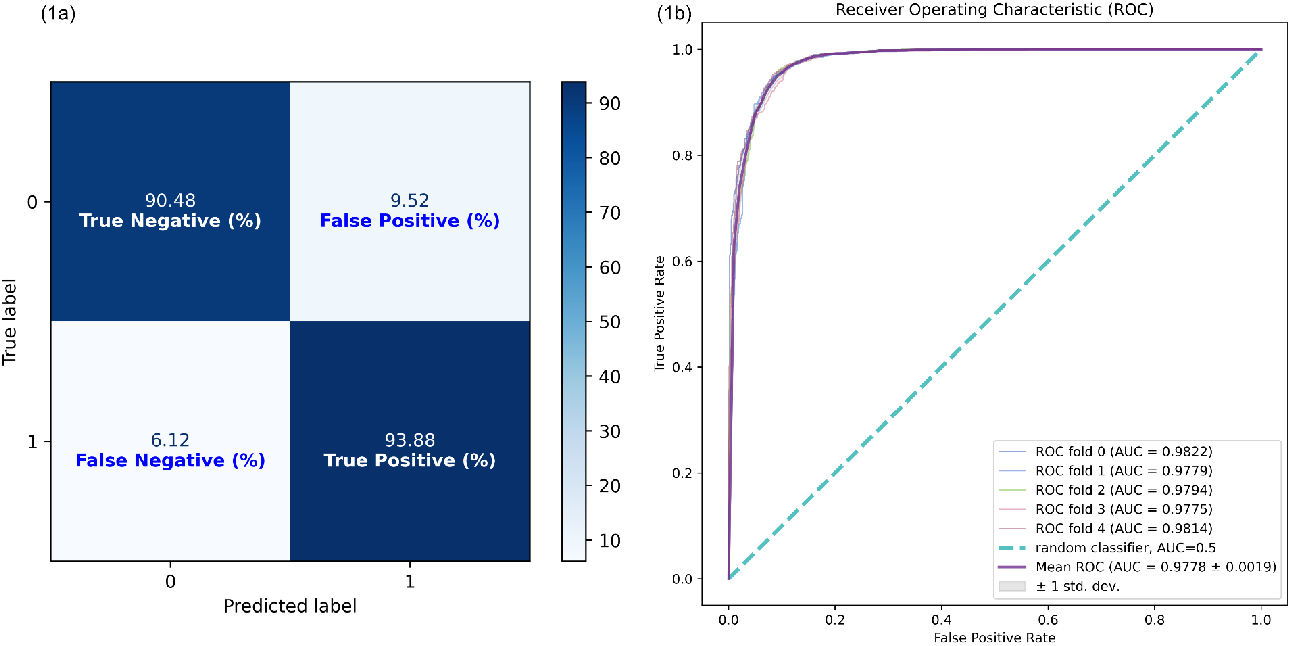
a) illustrating the confusion matrix and ROC curve for evaluating the RF model. The percentages of the confusion matrix display the proportions of true or false matches per class. For example, the model predicts 93.88% of true class 1 inputs. b) The ROC curve summarised the results of five cross-validation and the mean AUC (area under curve) is highlighted with one standard deviation.

### Identifying RF feature importance

After evaluation, we inspected the Gini importance of each feature provided by `Scikit-learn` (Supplementary Fig. 1). Gini importance calculates how a feature reduces impurity (based on the heterogeneity of node splits) across all trees in random forest models ^34,49^. We observe that MS2-based characteristics are the most important features. This indicates that MS2 data quality and ionisation efficiency affects the detection of deamidated peptides. Additionally, side-chain carbonyl carbon - backbone nitrogen distance and half-time estimates are the most significant physico-chemical features. These features are associated with the formation of ring intermediates (succinimides or glutarimides) via the nucleophilic attack of the side chain by the backbone nitrogen of the next amino acid (n+1). Half-life estimates are also based on primary sequences and neighbouring amino acids. It seems that Psi (ψ, an angle defining the alpha carbon and the carboxyl part) is more impactful than Phi (ϕ, an angle determining the rotation between the alpha carbon and the amino side) angle. This is consistent with the literature that the C-terminal of asparagine or glutamine is more involved in deamidation than the N-terminal ^11^.

We note that solvent exposure measurements have a limited impact on the model. Similarly, it is not significantly influenced by disorder predictions (IUPred scores). However, a caveat is that SASA (solvent accessible surface area) estimations are not adjusted by nearby amino acid conformation and orientation ^50^, which may affect solvent exposure at the residue level. Additionally, IUPred3 utilises an energy estimation matrix ^51^ that may not capture the full complexity of structural changes during degradation. Notably, as diagenesis progresses, BLG has the propensity to form amyloid fibrils driven by the collapse and aggregation of beta sheets during proteolysis, hydrolysis, or extreme pH conditions ^52–54^.

### Fast deamidation sites in modern, degraded BLG

We heated BLG and mixtures (BLG + pure glucose, BLG + octanoic acid (C_8_), and BLG + glucose + C_8_) at 70 °C, pH 7 from 0 to 12 hours to investigate the onset of deamidation. The inclusion of sugar and C_8_ acid is relevant. Dietary proteins may have been incorporated into archaeological materials via multiple and dynamic pathways, including cooking, chewing, and complex interactions between microbes and hosts ^36,55^. It is essential to investigate the dynamics between proteins, sugars, and lipids given their roles in protein modifications and preservation ^56,57^. C_8_ was selected based on its slight water solubility without any treatment ^58^. For BLG, temperatures between 65 °C and 70 °C at pH 7 are characterised by dissociating tertiary but mostly intact secondary structures, known as unfolding, melting globular states ^59,60^. However, above 70 °C, irreversible losses of secondary structures occur, especially for beta sheets, resulting in increased disordered structures and/or the formation of aggregates ^61,62^. Hence, the combination of 70 °C and different BLG matrices were chosen to monitor interactions between the starting point of irreversible conformational changes, complex mixtures, and deamidation kinetics.

We calculated position-specific deamidation abundance, adjusted by probabilities obtained from the RF model and MS2 intensities normalised for amino acids sharing a position. We used polynomial regression to calculate the rate of change, since a simple linear regression failed to fit the data points (Supplementary Fig. 2).

While the relationship between deamidation and time is not linear in degraded BLG, we observe clear fast and slow deamidation sites (Fig. 2). Indeed, the overall pattern is that asparagine deamidates more rapidly than glutamine. However, two outliers are identified: LQ[75]K and EN[106]K. LQ[75]K is the only position where fatty acid-containing BLG mixtures exhibit fast deamidation. One explanation is that this position is located within the beta-barrel. BLG is a lipocalin and its barrel contains binding sites for small, non-polar compounds, including retinols and fatty acids ^63^. It is also reported that the carboxyl end of a fatty acid interacts with charged residues (for example, lysine here after Q[75]) via nonspecific hydrogen bonding and electrostatic interactions ^64^. The beta-barrel is relatively open at pH 6-8 ^65^ and is dissociating when heated at 70 °C ^59,60^, which may have enabled the interactions between C_8_ and lysine at this position under the experimental conditions (70 °C, pH 7). However, a paradox arises because the C_8_-lysine complex is expected to be bulky, which may hinder, rather than accelerate, deamidation.

**Fig. 2.**
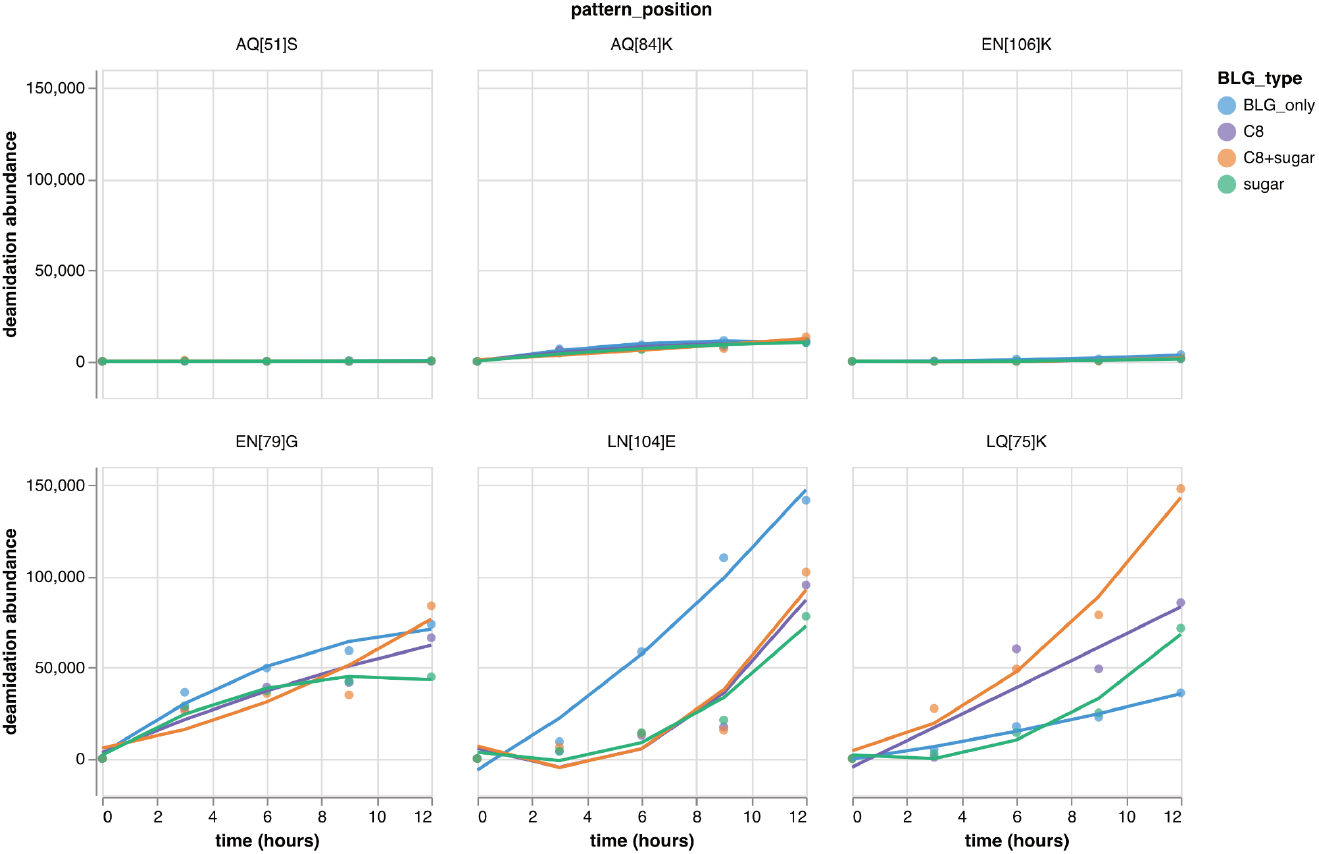
Polynomial regression plots illustrate slow (top row) and fast (bottom row) deamidation sites in modern, experimentally degraded BLG.

It is plausible that C_8_-lysine interactions alter the local conformation. While the carboxylate (COO^-^) of C_8_ interacts with the lysine positively charged side chain (NH3^+^), the aliphatic, non-polar hydrocarbon chains may move towards other hydrophobic amino acid side chains, especially tryptophan adjacent to lysine (LQ[75]KW) through hydrophobic interactions. This hydrophobic clustering may push and expose glutamine to water. Alternatively, the clustering and heat degradation of C_8_ over time may release volatile by-products, including aldehydes (R-CH=O) and/or free radicals. They may affect local entropy, or participate in Maillard interactions ^57,66^. However, it remains unclear how Maillard interactions and complex, multi-stage intermediates (Amidori products/ alpha-dicarbonyls) affect deamidation kinetics.

Additionally, we observe that EN[106]K overlaps with LN[104]E, but the latter exhibits faster deamidation rates. This is supported by sequence-based deamidation half-time estimates, where LN[104]E is associated with a deamidation half-life of 56.7 days ^67^. Additionally, this position also has a higher solvent accessible surface (58.33 square Å) than that of the EN[106]K (7.53 squared Å). However, there is no substantial difference in the nucleophilic attack distance between the two positions, and EN[106]K may still deamidate over time.

### A multi-level approach for ancient protein authentication

We used the same method to calculate position-specific, probability-adjusted, and intensity-normalised deamidation abundance for 102 MS2 files from three archaeological datasets ^38,42,43^. These datasets were selected because of the abundant BLG PSMs reported in the original publications. Fig. 3a shows that LQ[75]K is the most frequent and highly deamidated site. This indicates that fatty acids or other short, non-polar compounds may have been present in the archaeological matrices, binding to the beta-barrel and affecting glutamine deamidation at this position, consistent with the deamidation patterns observed in the degraded, fatty acid-containing BLG mixtures. We also identify that LN[104]E and EN[106]K are other two advanced deamidation sites. Notably, it is hypothesised that slow deamidating sites such as EN[106]K display advanced deamidation in archaeological samples given the extended time scale. However, the patterning is not consistent, as AQ[51]S, another slow deamidating site under experimental conditions, fails to exhibit high deamidation over time. Given the variability, it remains inconclusive whether stand-alone, position-specific deamidation patterns are sufficient to authenticate ancient proteins.

**Fig. 3.**
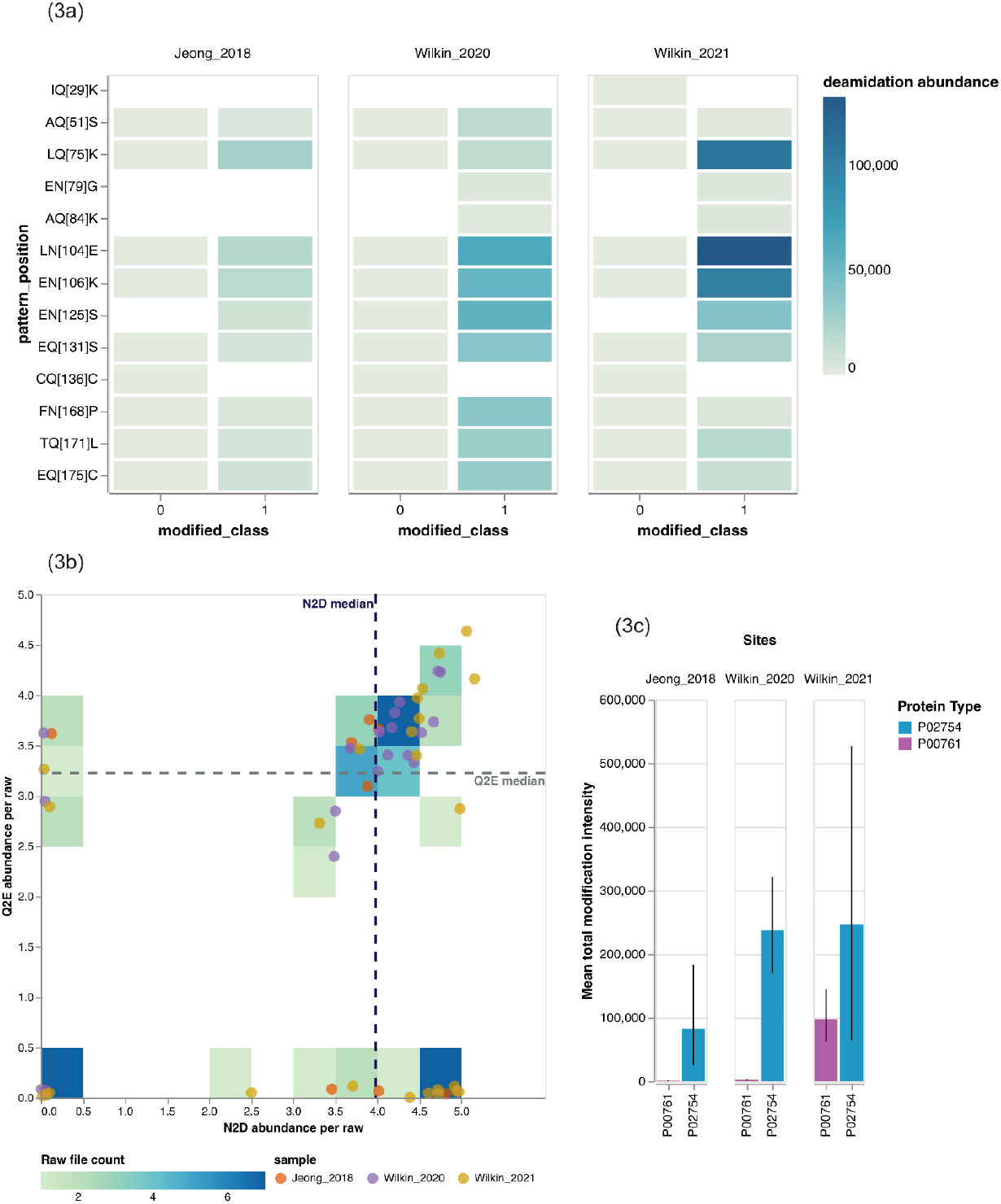
a) highlights frequent deamidation sites in the three ancient BLG datasets. b) demonstrates N2D vs Q2E clustering with medians. Abundance values were transformed by log_10_. c) illustrates overall and weighted variable modifications at the protein level comparing porcine trypsin (P00761) with BLG (P02754).

We also plotted asparagine (N2D) against glutamine (Q2E) deamidation abundance to see if putative, ancient BLG demonstrates high levels of asparagine and glutamine deamidation. While there are two clusters of no deamidation and only high asparagine deamidation, we observe that 33.33% of the archaeological raw files demonstrate advanced asparagine and glutamine deamidation above the medians (Fig. 3b). The presence of no deamidation and asparagine-only deamidation is problematic, since this indicates risks of contamination and deamidation introduced during extraction or other laboratory procedures. It is reported that types of buffers (especially those containing acids), heating, and incubation time may introduce laboratory artefacts and artificially accelerate deamidation ^68^.

Given the clustering results, we included an additional layer of authentication by comparing the overall modification levels of putative ancient BLG with trypsin processed in the same batch. We calculated weighted, normalised, and overall variable modification abundance for BLG and trypsin without the inclusion of RF probabilities. Trypsin was selected as a baseline, as it is widely used in bottom-up proteomics and is added to each sample for enzymatic digestion. We incorporated all variable modifications (including deamidation and oxidation of methionine) to account for BLG PSMs that do not contain glutamine or asparagine to minimise bias. Fig. 3c shows a clear trend that ancient BLG is more modified than trypsin. This is supported by non-parametric Mann-Whitney U tests that there are significant differences across all datasets (all p-values < 0.05, Supplementary Table 1).

Despite the clear protein-level distinction, we acknowledge that the use of trypsin only monitors deamidation introduced during and after digestion. Additionally, trypsin deamidation may be exacerbated by MS2 workflows. For example, trypsin digestion is often quenched by adding acids to lower the pH. Similarly, conventional sample cleaning up procedures such as stage tipping require samples to be acidified with formic acid or trifluoroacetic acid (TFA) before loading ^69^. In liquid chromatography, acids (formic, trifluoroacetic, or acetic acids) are often added to the mobile phase to increase sample solubility, promote protonation, reduce background noise, and facilitate separation using a (C_18_ silica-packed) stationary column ^70^. Hence, the widespread use of acids in MS2 analysis may inadvertently accelerate trypsin deamidation.

Nevertheless, these results underscore the importance of incorporating multiple lines of evidence to authenticate ancient proteins. The combination of position-specific patterns, N2D vs. Q2E clustering, and overall variable modifications at the protein level is useful to disentangle the variability and often context-specific deamidation behaviours. We leverage a high performance random forest classification model to incorporate a wide range of MS2-based and physico-chemical features and investigate their importance. While this model is BLG-focused, it opens up new possibilities for other archaeological substrates and model degradation experiments.

## Materials & Methods

### Ancient BLG datasets

Ancient BLG datasets were collected from PXD008217 ^38^, PXD014730 ^42^, and PXD022300 ^43^. These datasets were selected based on the rich BLG peptides reported in the original publications. Their .RAW files are also complete and publicly available.

### BLG degradation experiments

Purified BLG lyophilized powder (≥ 90%, CAS: 9045-23-2), pure glucose powder (≥ 99.5%, CAS:50-99-7), and pure octanoic acid (C_8_) in solution (≥ 99%, CAS: 124-07-2) were obtained from Sigma-Aldrich^®^. Four BLG-based degradation experiments were set up: BLG-only, BLG + glucose, BLG + C_8_, and BLG + glucose + C_8_.

For the BLG-only fraction, a BLG stock solution was diluted to 0.2 μg/μL in molecular biology grade water (BN-51100, BioNordika). The BLG concentration in all mixed fractions containing glucose and/or C_8_ was also maintained at 0.2 μg/μL. A 1:1 molar ratio of BLG to glucose or C_8_ was used for the binary combinations, and a 1:1:1 molar ratio was used for the BLG + glucose + C_8_ mixture. All solutions were pH 7 and confirmed by pH strips (MQuant^®^). Eppendorf^®^ Protein LoBind tubes were used for all the sample fractions to minimise sample binding and loss.

Method blanks consisting of molecular biology grade water, as well as samples were incubated at 70 °C in a heating block for a total of 12 hours. At 0, 3, 6, 9, and 12 hours, 100 μL was sampled from each of the four degradation conditions; this was done in triplicates for a total of 300 μL. All the aliquots were digested by 0.4 μg trypsin (Sequencing Grade Modified, Promega) at 37 °C for four hours.

### LC-MS/MS analysis

All degraded BLG fractions and the blanks were acidified using 10% v/v TFA to below pH 2. The acidified samples were loaded, concentrated, and desalted on Evotips (Evosep Biosystems) as per manufacturer’s instructions. Each Evotip also functioned as a single, disposable trap column for an Evosep one HPLC (high performance liquid chromatography) system (Evosep Biosystems). A pre-set Evosep 60 SPD method (60 samples per day) was used with an 8 cm EvoSep C_18_ Endurance column (EV1064, 8 cm x 100 μm, 3 μm beads).

The Evosep one system was coupled with a Q Exactive™ HF (high-field) hybrid mass spectrometer (Thermo Scientific™) with an electrospray ionisation source (ESI). Full scans (MS1) were acquired over a mass-to-charge (m/z) range of 350-1600 at a resolution of 120,000, with an automatic gain control (AGC) target value of 3E+6 and a maximum injection time (IT) of 100 ms. The 12 most intense precursor ions from charge state +2 to +4 were isolated within a window of 1.2 m/z and fragmented with a normalised collision energy (NCE) of 28. The dynamic exclusion was set to 1 s to temporarily exclude most-abundant precursors. MS2 spectra were acquired at a resolution of 30,000, with an AGC target value of 1E+5 and a maximum IT of 100 ms.

### MS2 Data conversion

All ancient and experimentally degraded BLG .RAW files were converted to vendor-free .mzML files by `ThermoRawFileParser` (1.7.3) using default parameters without file size reduction.

### Orthrus MS2 data analysis

All collected, ancient and acquired, degraded BLG MS2 spectra were analysed by `Orthrus` (v 1.0.0). Briefly, the non-enzymatic model provided by `Casanovo` was used for the ancient BLG data, while the tryptic model was applied to the degraded BLG spectra.

For `Sage` database searching, tryptic digestion without proline suppression was selected, with two missed cleavages. The carbamidomethylation of cysteine was set as a fixed modification. Variable modifications were deamidation (asparagine + glutamine) and the oxidation of methionine, with a total of five modifications allowed per identification. In accordance with `MaxQuant` optimised settings for Orbitrap™ instruments, the MS1 mass tolerance was 7 ppm, and the MS2 mass tolerance was 20 ppm.

A joint model was used for ancient and degraded BLG search results during `Mokapot` rescoring to mitigate the issues caused by low-abundance samples ^71^.

### RF features and implementation

**Methods Table 1:** 13 features selected for the RF classification and their measurements. All features are continuous (containing decimals).

| RF feature | Description | Source |
| --- | --- | --- |
| Attack distance | The distance (Å) between a side chain carbonyl carbon of asparagine or glutamine and the backbone nitrogen of the next amino acid (n+1). | Ordinates collected from a protein PDB file and calculated by Biopython. |
| Half-life estimates | Published half-life estimates of synthetic peptides (for example, XxxAsnYyy and XxxGlnYyy). | Synthetic peptides were kept at pH 7.4, 37 °C, 0.15 M Tris-HCl buffer for 300 days to estimate deamidation rates <sup>67</sup> |
| Psi angle ($\psi$) | An angle between the alpha carbon and the carboxyl group (CA-C). | Ordinates collected from a protein PDB file and calculated by Biopython. |
| Phi angle ($\phi$) | An angle between the alpha carbon and the amino group (CA-N). | Ordinates collected from a protein PDB file and calculated by Biopython. |
| SASA | Solvent accessible surface area, measuring the exposure of a residue to a solvent (squared Å). | Calculated by Shrake & Rupley algorithm as part of a Biopython module <sup>72</sup>. |
| Adjusted pI | Isoelectric point, the pH at which a peptide carries no charge. The calculated pH is adjusted by subtracting pH 7. | Calculated by Biopython. |
| IUPred | Disorder propensity. | Obtained from IUPred3 website tool <sup>51</sup>. |
| Tryptic ratio | A ratio of the length of an assigned peptide to its fully tryptic version. A fully peptide has a value of 100.0. | Python Pandas dataframe operation. |
| Adjusted FDR | A ratio of a reported FDR to the threshold 0.01. | Python Pandas dataframe operation. |
| Intensity ratio | A ratio of a reported MS2 intensity value to an overall intensity median. | Python Pandas dataframe operation. |
| Matched ion intensities (%) | The percentage of matched ion intensities between acquired and theoretical spectra. | Collected from a Sage output file <sup>73</sup>. |
| Retention time difference | The difference between observed and predicted retention time. | Collected from a Sage output file <sup>73</sup>. |
| Precursor difference (ppm) | Mass difference between an observed and theoretical precursor mass. | Collected from a Sage output file <sup>73</sup>. |

The protein structure file of BLG was fetched from RCSB Protein Data Bank (identifier: 1BEB, resolution: 1.8 Å). As described in Methods Table 1, the 1BEB. pdb file was used to calculate the attack distance, dihedral angles, and SASA by `Biopython` (1.84). Published half-time estimations ^67^ of asparagine and glutamine synthetic peptides were included. Disorder propensity was determined by IUPred3 as part of their interactive web server ^51^, and the sequence of P02754 (UniProt identifier for bovine BLG) was used as the input. MS2-based features were collected from `Sage` (0.14.6) output files (results.sage.tsv). Random forest binary classification was implemented using `Scikit-learn` (1.5.2) on Colab.

### RF hyperparameter fine tuning

We used random search (RandomizedSearchCV, `Scikit-learn`) to gauge the hyperparameter ranges, and subsequently applied grid search (GridSearchCV, `Scikit-learn`) for precise tuning. Hyperparameter tuned were reported in Methods Table 2. ROC-AUC analysis was used to evaluate the default `Scikit-learn`, tuned random search, and best grid search models.

**Method Table 2:** Hyperparameter ranges for the random search. The narrowed-down hyperparameter space was refined by grid search.

| RF hyperparameter | Description | Range |
| --- | --- | --- |
| n_estimators | The number of trees in the forest. | randint(50,500), randomly selected between 50 and 500 |
| max_depth | The maximum depth of each tree (layers) | [None, 10, 20, 30] |
| min_samples_split | The minimum numbers of samples required to split an internal node. | [2, 5, 10] |
| min_samples_leaf | The minimum numbers of samples required to split an external node. | [1, 2, 4] |
| max_features | The number of features to consider when splitting at each tree node.<br>sqrt= the square root of the total features.<br>log2=the log<sub>2</sub> transformation of the total features. | ['sqrt', 'log2'] |

### RF evaluation

The effectiveness of the RF binary classification was evaluated by precision, F1 scoring, a confusion matrix and ROC-AUC analysis with 5-fold cross validation. All evaluation matrices were obtained from `Scikit-learn` (1.5.2).

### Position-specific deamidation estimations

Deamidation abundance was calculated at the position level. Each abundance value was based on the MS2 intensity of a PSM, normalised by each amino acid constituent and adjusted by a RF prediction probability. For example, if a .RAW file contains two BLG peptides: TKIPAVFKIDALN[+0.9848]EN[+0.9848]K (MS2 intensity per PSM=2324001, normalised intensity per amino acid=145250.06, RF prediction probability=0.64) and TKIPAVFKIDALN[+0.9848]ENK (MS2 intensity per PSM=1086790, normalised intensity per amino acid=67924.38, RF prediction probability=0.82), the deamidation abundance at LNK is 148658.03, and 92960.04 at ENK.

### Protein-level modification measurements

Since the RF model was BLG-focused, we included another level of measurement for general purpose without the adjustment of RF prediction probabilities. All variable modification abundance, including oxidation (methionine) and deamidation (asparagine + glutamine). Glutamine deamidation was weighted using the difference between glutamine and asparagine halt-life estimations ^67^ to reflect its time-dependent nature. The overall modification abundance of trypsin was used as a baseline.

### Data & code availability

Anubis is an open source project. Tutorials and pre-configured Jupyter Notebooks are available on Github (https://github.com/yc386/anubis_palaeoproteomics). The Github repository includes the BLG-focused RF model, the training datasets, and all Jupyter Notebooks used in the study for data analysis/figures.

### Grant Information

This project has received funding from the European Union’s Horizon 2020 research and innovation programme under the Marie Skłodowska-Curie grant agreement No. 956351. MJC acknowledges support from Danmarks Grundforskningsfond (DNRF128) and Carlsbergfondet (CF18-1110).

## Supplementary information

**Supplementary Fig 1:**
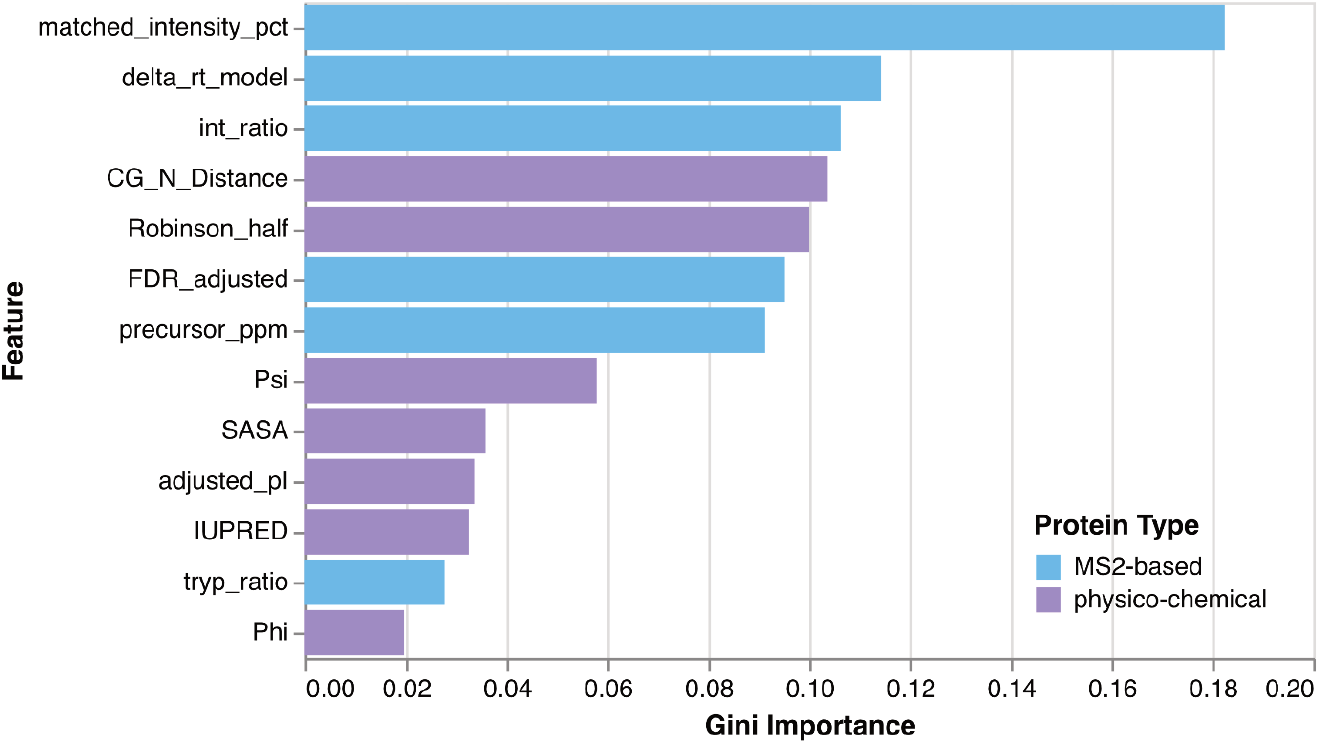
Gini importance values were obtained from `Scikit-learn` (1.5.2). MS2-based features were coloured in blue and physico-chemical characteristics in purple. Overall, matched_intensity_pct (% intensities matched between acquired and theoretical ion series), delta_rt_model (difference between observed and predicted retention time), and int_ratio (MS2 intensities normalised by intensity median) are among the most important features. Similarly, CG_N_Distance (distance between the side-chain carbonyl of asparagine/glutamine and the backbone nitrogen of the next amino acid), Robinson_half (published half-life estimations), and Psi (angle between the alpha carbon and the carboxyl group) are the most significant physico-chemical features. However, a caveat is that gini importance is biassed towards continuous values with a large number of potential split points ^74^. In the context of this BLG-focused model, matched_intensity_pct had possible 23269 split points, while CG_N_Distance had 13 potential splits.

**Supplementary Fig 2:**
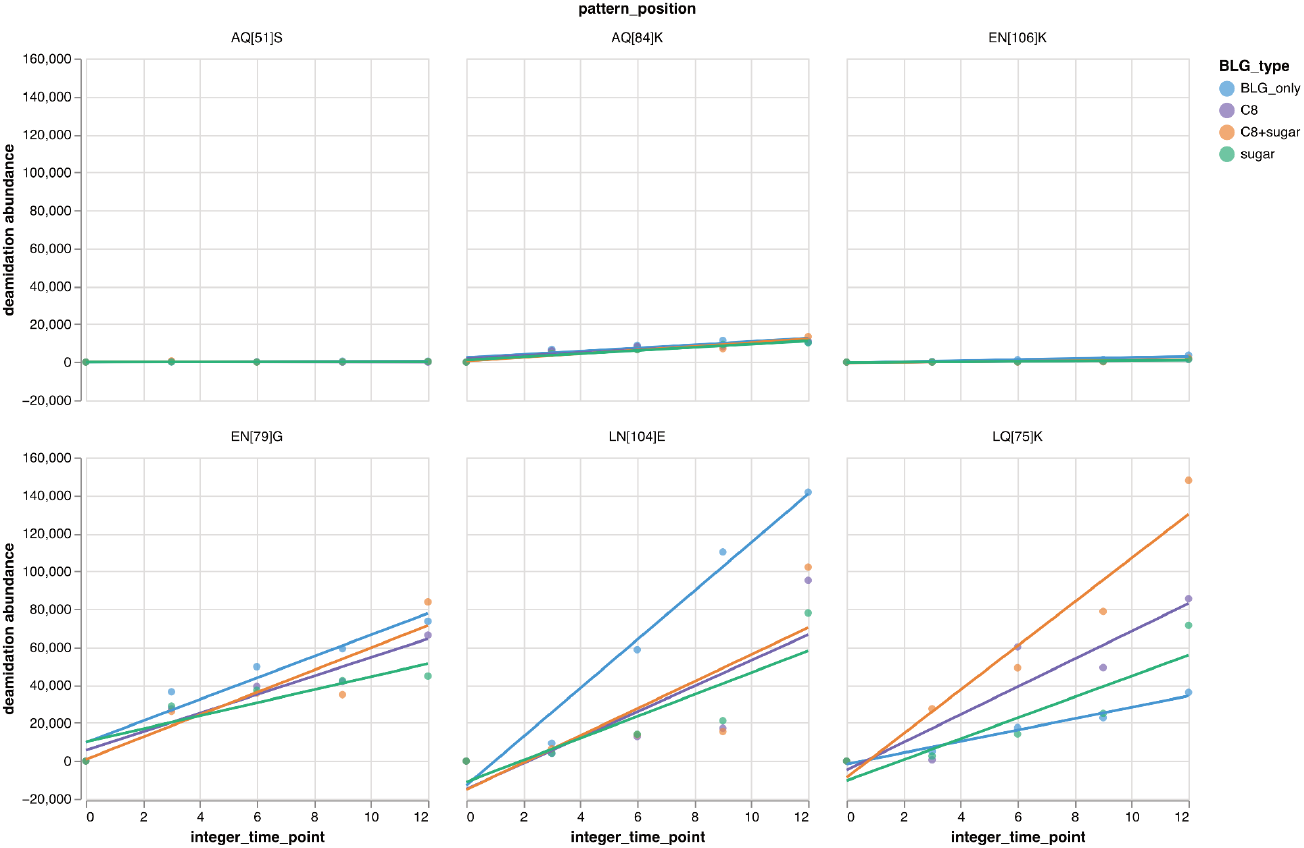
Linear regression (`Scipy` 1.14.1) was used to fit deamidation abundance over time. However, linear correlation was not clear and R-Squared (the coefficient of determination) values were low for some positions.

**Supplementary Table 1:** Mann–Whitney U tests (Scipy 1.14.1) were used to compare the deamidation abundance of putative ancient BLG and trypsin processed in the same batch. The *p* value was below 0.05 for each dataset.

| Putative ancient BLG vs modern trypsin deamidation abundance per dataset |  |  |
| --- | --- | --- |
| PRIDE accession number | Mann–Whitney U test statistic | *p* value |
| **PXD008217** | 94191 | 1.32E-19 |
| **PXD014730** | 520590 | 1.88E-82 |
| **PXD022300** | 652113 | 0.01 |

